# Photovoltaic enzymes by design and evolution

**DOI:** 10.1101/2022.12.20.521207

**Authors:** H. Adrian Bunzel, James A. Smith, Thomas A. A. Oliver, Michael R. Jones, Adrian J. Mulholland, J. L. Ross Anderson

**Affiliations:** School of Biochemistry, University of Bristol, University Walk, Bristol BS8 1TD, UK; Centre of Computational Chemistry, School of Chemistry, University of Bristol, Bristol BS8 1TS, UK; Department of Biosystems Science and Engineering, ETH Zurich, CH-4058 Basel, Switzerland; School of Chemistry, Cantock’s Close, University of Bristol, Bristol BS8 1TS, UK

## Abstract

The global energy crisis challenges us to develop more efficient strategies for the sustainable production of energy. Given the excellent efficiency of the natural photosynthetic apparatus, biohybrid photovoltaic devices present an attractive solution for solar energy conversion. However, their composition, stability, and complexity can limit their inclusion into photovoltaic devices. Here, we combined computational design and directed evolution to overcome these limitations and create tailor-made photoenzymes. Photo-biocatalysts were designed by introducing photosensitizer binding sites into heme-containing helical bundle proteins. The designed binding sites were specific for the target photosensitizer and readily transplanted into other helical bundles. The best design was highly evolvable and reached nanomolar ligand affinity after mutagenesis and screening. The evolved enzyme generated 2.6 times higher photocurrents than the photosensitizer alone, primarily driven by increased photostability. Evolvability is a unique advantage of our protein-based approach over abiological photovoltaic and will be critical to developing efficient biohybrid systems.

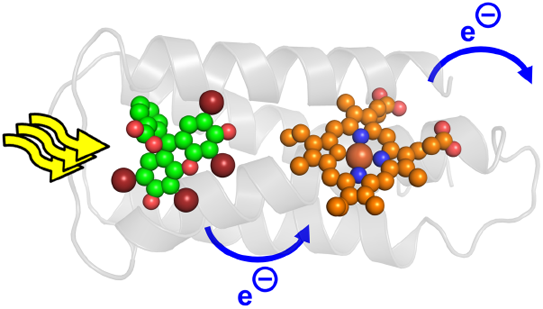

---

We are experiencing a global energy crisis that demands innovative and efficient strategies for sustainable energy production. To that end, photosynthesis, nature’s solution for solar energy conversion, can provide inspiration. Natural photosystems accomplish light harvesting with nearly perfect quantum yields.^1,2^ Exploiting these sophisticated biological complexes in biohybrid devices has been a long-sought-after goal in bioengineering.^3^ However, using natural photosystems in photovoltaic applications has proven difficult, partly because of poor protein stability and lack of control over the electron flow. Biohybrid solar cell efficiencies have thus fallen short of expectation,^3^ trailing well behind established silicon (26%),^4,5^ perovskite (24%)^6^, and dye-sensitized (13%)^7^ solar cells.

*De novo* photoactive proteins could potentially circumvent many limitations of natural photosynthetic enzymes. For instance, helical bundle proteins have been rationally designed that display some photoactivity after binding small-molecule photosensitizers. These systems typically accomplish photosensitizer-binding by bioconjugation or metal coordination^8–10^. This approach, however, is restricted to a limited set of photosensitizers and frequently yields ill-defined protein-cofactor interactions that compromise photoactivity. Instead of using solely rational methods to design proteins, computational protein design may provide a more accurate and general path to *de novo* photoenzymes. Nowadays, *de novo* proteins^11,12^ and simple enzymes^13^ can be computationally designed from scratch. While proteins that bind small-molecule dyes have been designed in the past,^14^ this approach has, to the best of our knowledge, not yet been used to design photosensitizer binding packets to create novel photoenzymes.

Regarding photo-biocatalysis, it is worth noting that many redox enzymes already display modest, non-specific photoactivity when mixed with photosensitizers in solution.^15^ However, the achieved efficiencies are often low, and improving activity to practically useful levels is challenging. In contrast, computationally designed enzymes can be systematically optimized by directed evolution.^16,17^ In fact, computationally designed enzymes are typically highly amenable to evolution, and several designs have been improved by orders of magnitude *via* rounds of mutagenesis and screening.^18–21^ We hypothesize that *de novo* photo-biocatalysts containing computationally designed photosensitizer binding pockets may likewise be evolvable to efficient made-to-order photoenzymes (**Fig. 1**).

**Fig. 1.**
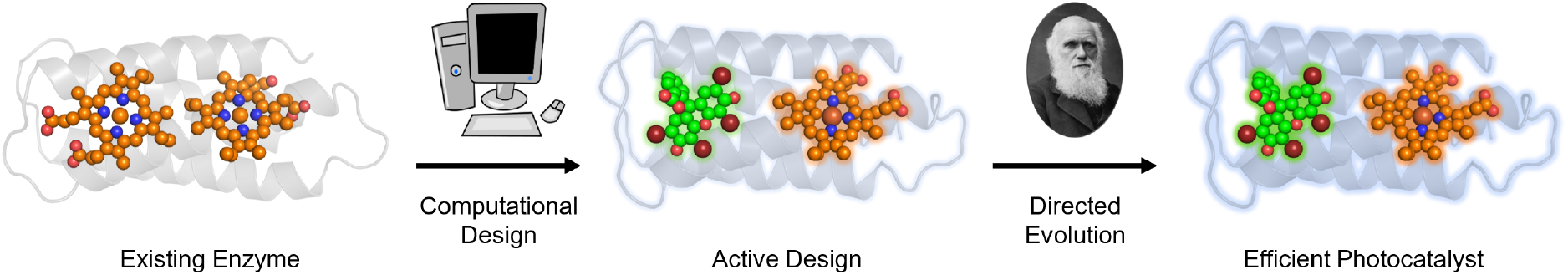
Creating photocatalytic enzymes by design and evolution. Computational design and directed evolution of photosensitizer binding sites affords photocatalytic activity.

Here, we created a photovoltaic enzyme by computational design of a photosensitizer binding site into a heme-containing protein, and subsequently improved photosensitizer affinity by evolution (**Fig. 1**). The best-performing variant was more photostable and generated 2.6 times more photocurrents than the photosensitizer alone. We anticipate that the inherent evolvability of our system could provide a competitive advantage over established abiological photovoltaic technologies that cannot benefit from directed evolution.

## RESULTS

### Computational Design

To create photovoltaic enzymes, we computationally designed photosensitizer binding sites into heme-containing proteins (Fig. 2a). Our design approach combined Rosetta^22–25^ with molecular dynamics (MD) simulations^26,27^ and was based on the helical bundle protein 4D2 that binds two heme cofactors.^28^ Design was tasked with replacing one of 4D2’s heme cofactors with the photosensitizer eosin Y (EOY).^29^ To generate conformational diversity in 4D2, 100 ns MD simulations were performed prior to design. Next, RosettaMatch^22^ was used to screen the resulting protein backbone conformations for suitable positions that fit EOY. Initially developed for enzyme design, RosettaMatch requires the definition of catalytic interactions to identify potential ligand binding sites. Here, the matcher was tasked with proposing EOY binding poses that can realize two H-bonding interactions with either of the ligand’s oxyanions. RosettaMatch afforded ≈10^3^ binding modes. For each identified binding mode, ≈100 EOY-binding variants were designed using Rosetta FastDesign.^23–25^ All designs were subsequently ranked by their overall score, number of H-bonds to the ligand, EOY solvent-accessible surface area, and distance between EOY and heme. 200 potential hits were identified that had snug ligand binding pockets with good shape complementarity to EOY, and EOY-heme distances <10 Å to facilitate rapid electron transfer.^30,31^

**Fig. 2.**
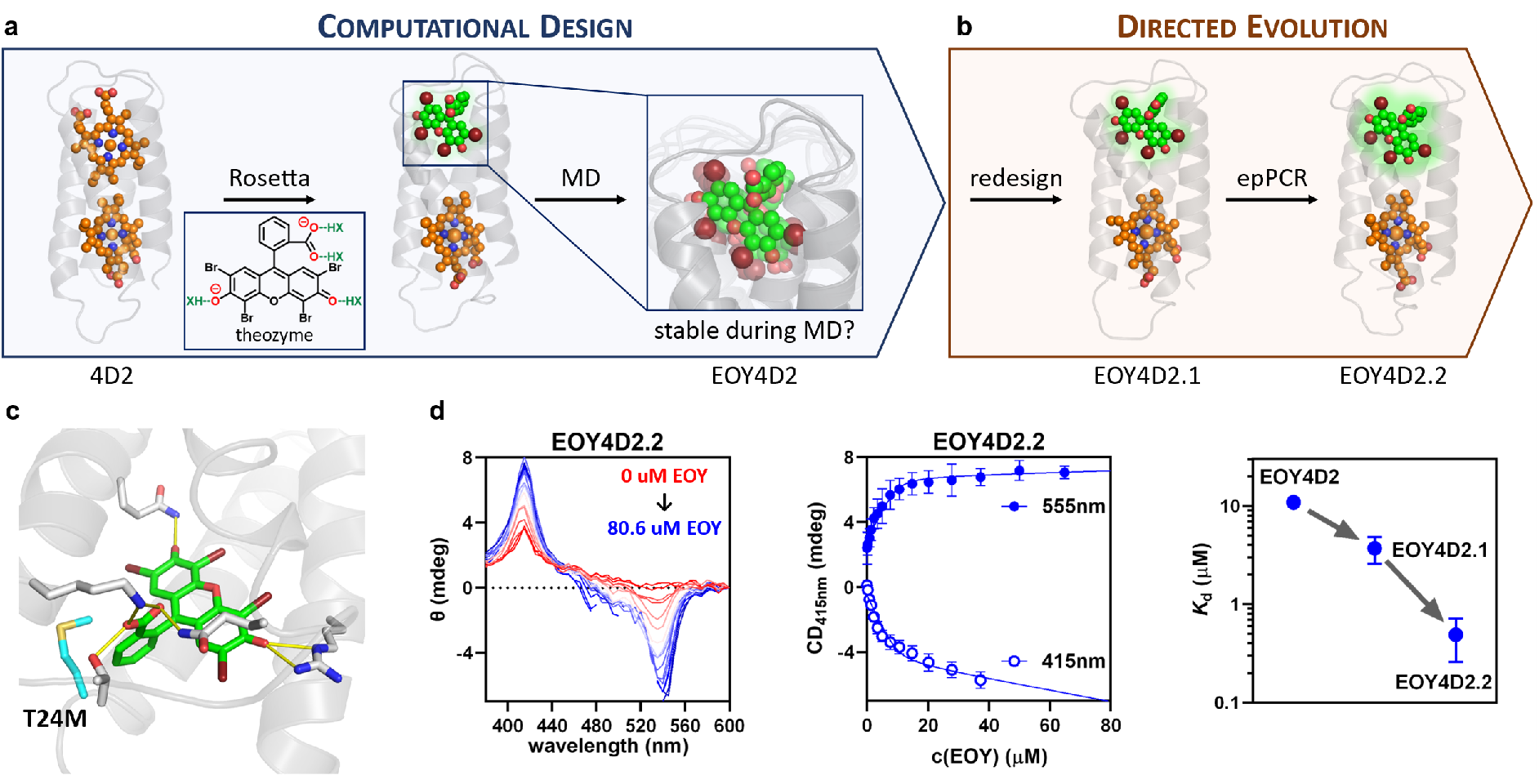
Design and evolution. **a**, Rosetta (match and design) was combined with MD simulations to design EOY4D2 computationally. RosettaMatch introduced EOY hydrogen-bonded to at least two protein residues (inset, green) into the protein scaffold 4D2. **b**, The protein was improved in two rounds of directed evolution. The first round was based on computational redesign, and the second on error-prone PCR (epPCR), resulting in EOY4D2.2. **c**, Modelling of EOY4D2.2 reveals distinct H-bonding interactions that probably lead to selective EOY recognition. (green: EOY, grey: protein, yellow: H-bonds, cyan: T24M mutation discovered after epPCR). **d**, CD titration showed that evolution increased EOY affinity by 22-fold, resulting in a *K*_d_ of 490 ± 230 nM.

Traditionally, these designs would be tested experimentally. To increase the success chances of our design approach, additional *in silico* screening was performed with the 200 potential hits. 12 ns of MD simulation were run for each design. The MD simulations were used to test if the designed binding site was maintained over time and to estimate binding energies. The designs were ranked by how much EOY was displaced from its designed position, the average number of EOY H-bonding interactions with the protein, and the EOY solvent-accessible surface area. In addition, binding affinities were estimated from the MD trajectories using MM-GBSA.^27^ We experimentally tested the two highest-ranking hits after *in silico* screening. The highest-ranking design did not bind heme – a feature we did not test beforehand. However, the second-best design, EOY4D2, could be loaded with heme and bound EOY with an affinity of 10.9 ± 1.5 μM, as determined by circular dichroism (CD) titrations.

### Alternative Scaffolds

To demonstrate the universality of the design approach, we targeted an alternative helical bundle protein with our computational pipeline. The alternative design was based on C45, a rationally engineered heme-peroxidase.^32^ In contrast to the 4D2-based variants, this design solely relied on a computational model – no crystal structure has yet been reported for C45. Despite the absence of experimental structural data, one out of six tested designs successfully bound EOY with a *K*_d_ of 57 ± 5 μM as shown by ITC (Fig. S1g+h), demonstrating the general applicability of our approach.

### Directed evolution

We subsequently set out to further enhance binding in a proof-of-concept directed evolution experiment, during which we created libraries either by computational redesign or random mutagenesis (Fig. 2b). Redesigning EOY4D2 with our design protocol and experimentally screening six variants resulted in EOY4D2.1, which bound EOY 3-fold tighter than EOY4D2 (*K*_d_ = 3.7 ± 1.1 μM). Subsequently, an error-prone PCR library was created and screened for EOY binding, relying on fluorescence polarization to assay the amount of protein-bound ligand in cell lysate (Fig. S1f). Screening of ≈400 variants allowed us to identify the T24M mutation, which enhances the packing of the EOY binding site, as indicated by modeling using Rosetta and MD (Fig. 2c). Directed evolution – using computational redesign and random mutagenesis to create diversity – resulted in EOY4D2.2 that binds EOY with an affinity of 490 ± 230 nM, 22-fold tighter than the original design.

### EOY binding is specific and modular

We confirmed EOY binding to EOY4D2.2 using calorimetric (ITC) and spectroscopic (absorbance, fluorescence, fluorescence polarization, CD) methods (Fig. S1b-e). Because the ligand and enzyme are both colored, some spectroscopic methods were limited by interference and inner-filter effects. In contrast, ITC relies solely on thermal readouts and unambiguously confirmed EOY binding. ITC, however, requires large amounts of purified proteins and cannot provide the throughput required for directed evolution. Fluorescence polarization was thus used to screen for binding in cell-lysate and microtiter plates (Fig. S1f). CD, which is more accurate than fluorescence polarization but challenging to be performed in microtiter plates, was then used to quantify binding with purified enzymes.

CD titrations showed two EOY binding signals that stemmed from the heme and EOY ellipticity (θ_max_: 415 nm and 555 nm, Fig. 2d). Interestingly, the EOY ellipticity showed a second low-affinity binding event, which was absent for the heme signal. This second binding event suggests a weak promiscuous interaction of the dye with the protein, which was also observed by ITC (Fig. S1b). This weak promiscuous interaction could potentially give rise to non-specific photoreduction, which agrees well with the photoactivity of many redox proteins when mixed with EOY in solution.^15^ The increase in heme ellipticity probably is not caused by an increase in protein stability upon EOY binding, as the addition of EOY did not significantly affect the protein CD spectrum or thermostability of EOY4D2.2 (Fig. S2). Instead, the oppositely signed CD signals of EOY and heme indicate excitonic coupling, which might cause the observed increase in heme ellipticity.^33,34^ Long-range excitonic coupling indicates dipolar interactions between heme and EOY. Targeting these interactions by design and evolution may allow tuning the energy landscape of the photoinduced electron transfer to boost catalysis.

Optically transparent thin-layer electrochemistry was used to interrogate how EOY binding affects the heme electronics in EOY4D2.2 (Fig. S3). A slight decrease in heme midpoint potential was observed upon EOY binding (-141.3 ± 1.2 mV to -148.8 ± 1.9 mV vs. NHE), which would be expected for the negatively charged EOY stabilizing the heme Fe(III) state rendering it less reducing. Modulating the heme and EOY potential, either by computational design or directed evolution, would provide a strategy to improve the electron transfer efficiency in future variants.

In contrast to the weak non-specific interaction of EOY with the protein scaffold, the designed binding site seemed highly specific (Fig. 3a). None of the designs bound fluorescein, the non-brominated analog of EOY. Furthermore, EOY4D2.2 binds EOY 5-fold tighter than erythrosin B, in which the bromine atoms of EOY are replaced with iodine. The specificity for EOY indicates the successful creation of a binding pocket with excellent shape complementarity and suggests that computation predicted the correct EOY binding mode.

**Fig. 3.**
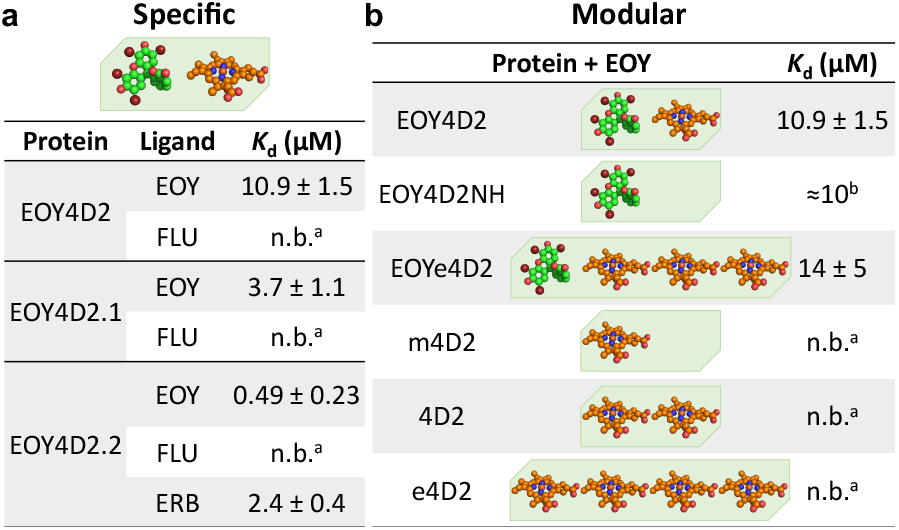
The designs are specific and modular. **a**, The EOY4D2 family selectively binds EOY over the structurally analogous fluorescein (FLU) and erythrosin B (ERB). **b**, The designed binding site is modular and can be readily transplanted in other helical bundle proteins (_a_ n.b. = no binding observed by ITC or CD titrations. _b_ *K*_d_ from ITC measurements).

EOY4D2 is furthermore highly modular (Fig. 3b). Without redesign, the EOY binding sequence in EOY4D2 was readily transplanted onto the helical bundle protein m4D2 (one heme) or e4D2 (four hemes), giving rise to constructs with one EOY binding site and none or three hemes (EOY4D2NH and EOYe4D2). This modularity will be advantageous in the future and could potentially allow combining modules with various functions in multi-cofactor systems. For instance, photosensitizer binding sites may be mixed and matched to tune the absorbance of the final catalyst mix to the inbound light.

### Photovoltaic activity

Having established that EOY tightly binds to EOY4D2.2, we next probed the protein’s photovoltaic activity in a model solar cell (Fig. 4). To that end, we assayed the short-circuit photocurrents generated from EOY4D2.2 and EOY in solution. To assay the photocurrents, samples were loaded onto screen-printed carbon electrodes and excited with a green LED (530 nm) while applying a potential of 0 V vs. NHE. Notably, EOY alone showed pronounced photobleaching, while the photocurrents measured for the photovoltaic complex remained relatively constant during the ≥8 h experiments. This improved photostability resulted in (2.6 ± 0.2)-fold increased photocurrents of EOY4D2.2 compared to EOY after prolonged excitation. The fact that EOY4D2.2 outperformed EOY after only two rounds of optimization promises that further evolution of the photoenzyme might result in highly efficient photo-biocatalysts for use in biohybrid solar cells.

**Fig. 4.**
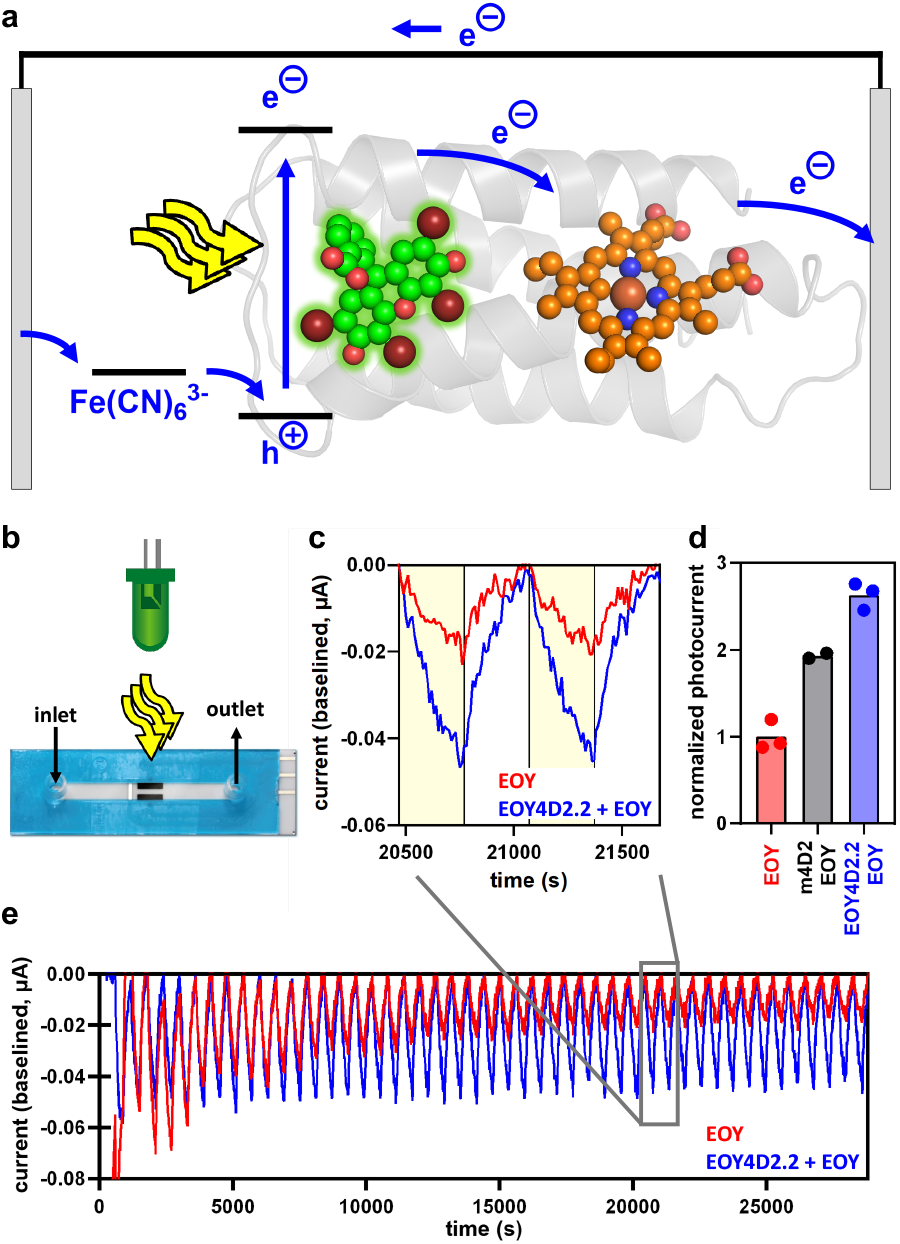
Photovoltaic activity. **a+b**, Photocurrents generated by EOY4D2.2 were assayed in solution using flow-cell electrodes after excitation with a green LED (530 nm). **c**, Photocurrents of EOY4D2.2 with EOY (blue) were higher than those of EOY alone (red). Yellow shades indicate when the LED was switched on. The baselining method is detailed in Fig. S4. **d**, EOY4D2.2 increases the photocurrents by 2.6-fold compared to EOY. Photocurrents were normalized to EOY alone and obtained from the area under the baselined curve. **e**, Prolonged excitation revealed that EOY4D2.2 boosts photovoltaic activity by preventing photobleaching.

## DISCUSSION

We have created a photovoltaic enzyme by computational design and directed evolution of a photosensitizer binding site into a heme-containing protein. The high photostability of our photoenzyme resulted in 2.6 times higher photocurrents compared to the photosensitizer alone. Taking inspiration from existing solar cells, we envision that our protein engineering approach may be used to amend existing dye-sensitized solar cells by packing the photocatalytic dye into a protein scaffold. Such organization might result in more efficient devices in which the evolvable protein provides tight control over the photo-electrochemistry and spatial organization of all redox partners.

Our photoenzyme is based on a helical bundle scaffold that is highly versatile and thermostable. The modularity will likely allow mixing and matching modules to combine various functions. Self-assembly of the photosensitizer with our designer catalysts – driven by a highly specific binding pocket – provides tight control over the system. The excellent shape-complementarity of the designed binding site omits the need to covalently link the photosensitizer with the protein, which will likely facilitate future *in vivo* applications. Most importantly, a defined binding site is essential to evolvability, as evolution is fundamentally based on tuning molecular interactions. We expect that further directed evolution will tailor the redox potentials and reorganization energies of all involved redox partners to boost photovoltaic activity while preventing deleterious side reactions. The inherent evolvability of our systems could potentially provide a competitive advantage over established abiological photovoltaic technologies that cannot benefit from directed evolution.

Our proof-of-concept work provides a valuable addition to the growing interest in accessing novel reactivities through the creation and evolution of photoenzymes.^35,36^ In particular, EOY can give rise to a diverse range of photochemistries^29^ that could be exploited in the future. We have demonstrated a methodological framework for enzyme photosensitization by design and evolution, which could potentially be used to create highly efficient photo-biocatalysts for various societal critical reactions. Computational design may be used to photosensitize virtually any redox enzyme, and subsequent directed evolution provides a methodological and systematic way to improve activity to practically useful levels. Thus, future design and evolution may allow targeting countless essential transformations – such as nitrogen fixation, carbon capture, and hydrogen production – with sustainable tailor-made photoenzymes.

## AUTHOR INFORMATION

We encourage critical feedback and suggestions on our preprint. Please submit any comments to.

## Author contributions

H.A.B. devised and performed the experiments. J.L.R.A. supported the experimental work, A.J.M. the computational work, and J.S., T.A.A.O. and M.R.J. the photocatalytic experiments. H.A.B. wrote the manuscript with input from all co-authors.

## Acknowledgments

HAB and AJM thank EPSRC (EP/M013219/1 and EP/M022609/1) and with JRA BBSRC (BB/M000354/1) for funding. HAB thanks the SNSF (postdoc.mobility, return, and ambizione fellowship) for support. TAAO acknowledges financial support from the Royal Society (URF\R\201007). This work is part of a project that has received funding from the European Research Council under the European Horizon 2020 research and innovation programme (PREDACTED Advanced Grant Agreement no. 101021207) to A.J.M. This work was conducted using the computational facilities of the Advanced Computing Research Centre, University of Bristol. We thank Martin Held for his helpful discussions on our manuscript.

## MATERIALS AND METHODS

### Computational Design

The computational design was performed in a three-step process comprising initial MD to generate conformational diversity within the scaffold protein, design using Rosetta Match,^22^ Fastrelax,^23^ and Fastdesign,^24,25^, and a final *in silico* screening step using MD. All MD simulations were run with Amber16 (sander.MPI for minimization and pmemd.cuda for MD simulations)^26^ on the University of Bristol HPC clusters (Bluecrystal Phase 3 and 4; BluePebble)

### Initial MD

MD simulations were performed similarly as described before.^28,32^ For the 4D2-based designs (PDB: 7AH0), the Amber ff14SB forcefield was used to parameterize the protein. Parameters derived for the bis-histidine ligated *b*-type heme in cytochrome *c* oxidase^37^ were used for heme, and GAFF-derived parameters were used for EOY. The system was built using tleap.^26^ Enzymes were solvated in an octahedral box of TIP3P water with a minimal distance of the protein to the box boundary of 8.0 Å. The net charge was neutralized by addition of sodium ions. For the C45-based designs, the systems were built with VMD. The CHARMM22 forcefield was used to parametrize the protein, *c*-type heme parameters from Autenrieth *et al*.^38^ were used for the heme, and CGenFF-derived parameters were used for EOY. The system was solvated in an 18 Å water box, 0.15 M NaCl was added, and neutralized with Na^+^ ions.

During minimization and equilibration, weak positional restraints were applied to all protein and cofactor atoms with a restraint weight of 10 kcal/mol/Å^2^. Langevin dynamics were used with a collision frequency of 0.2 and a 2 fs time step. All bonds involving hydrogens were constrained using the SHAKE algorithm. The systems were minimized using 500 steps of steepest descent followed by 500 steps of conjugate gradient minimization. Subsequently, the systems were heated from 0.1 K to 300 K in 50 ps in the NVT ensemble. After 50 ps MD in the NPT ensemble, the positional restraints were removed, and production was started in the NPT ensemble. For the initial 4D2-based design, snapshots were taken after 50, 100, 150, 200, and 300 ns and used as input. For the 4D2-based redesign, five 100 ns MD simulations were performed with EOY4D2.1 bound to EOY, and snapshots were taken every 5 ns for design, totaling 100 design input structures. For the C45-based design, snapshots at 50, 60, 70, 80, 90, and 100 ns were used. In general, supplying Rosetta with a more diverse set of input structures seemed to have improved the sampling of the fitness landscape and sequence diversity.

### Rosetta Design

Before design, the MD snapshots were relaxed using Rosetta FastRelax.^23^ Subsequently, EOY was matched into these snapshots.^22^ To that end, a ligand grid was prepared with the gen_lig_grids application. The grid was built around the heme that was replaced during the design. For the C45-based design, a dummy EOY molecule was placed manually between the four helices to guide the grid towards the non-heme containing half of C45. Based on these grids, Rosetta match was run using constraint files that included 2 H-bonding interactions to any of the oxyanionic groups in EOY. Subsequently, structures were optimized with Rosetta Fast Design^24,25^, typically generating 100 designs for each matched structure.

### *In silico* screening by MD

The MD simulations for *in silico* screening were performed analogously to the initial MD simulations described above. To screen hundreds of designs, only 2 ns of NPT equilibration and 10 ns of NPT production were performed per variant. After aligning the protein backbone to the initial design structure, the EOY RMSD was calculated. Designs in which the RMSD exceeded 3 Å on average during the final 2 ns of MD were considered failures. For the resulting designs, the average number of H-bonds and solvent-accessible surface area were determined. Furthermore, ligand-binding strengths were estimated using MM-GBSA as implemented in Amber.^26^ The final variants for experimental testing were selected based on the most promising combination of metrics and after visual inspection of the MD trajectories.

### Directed evolution

#### Library generation

Unless indicated, designs were expressed from a pET-21(+) vector (*Thermo Fisher*) with an N-terminal 6xHis tag and TEV cleavage sequence. Genes were commercially synthesized and cloned into the target vectors by *Twist* and *Eurofins*. Cloned vectors were transformed into Stellar cells (*Takara*) before plasmid purification using a miniprep kit (*NEB*). Error-prone PCR was performed using the GeneMorph II EZClone Domain Mutagenesis kit (*Agilent*) according to the manufacturer’s protocol. The DNA template for mega primer generation was amplified using Q5 DNA polymerase (*NEB*) and the T7 and T7term primer from EOY4D2.1@pet21(+). Megaprimers were then produced using the T7 and T7term primer with either 100 ng or 10 ng template to create two libraries with varying mutational rates.

#### Screening

The constructed DNA libraries were screened in 96-well plate format. After transformation into T7 Express *E. coli* cells (*NEB*), variants were picked into 2 ml deep-well plates containing 1 ml LB autoinduction medium (*Formedium*). The plates were sealed with semipermeable Breathe-Easier membranes (*Merck*), and cells were grown in a plate shaker for three days (1,000 rpm, 30°C). Cells were harvested by centrifugation (2,000 rpm, 10 min) and resuspended in 600 ul assay buffer (20 mM sodium phosphate, 20 mM sodium chloride, pH 8) with lysozyme, DNase I, and polymyxin B. Cells were lysed in three freeze-thaw cycles by freezing the plates for more than >2 h at -20°C and thawing for 30 min in a water bath at room temperature. The lysate was cleared by centrifugation (2,000 rpm, 10 min), and 200 ul lysate was mixed with 50 ul reagent solution containing 25 uM hemin (final concentration 5 uM) and 25 uM EOY (final concentration 5 uM) in assay buffer (20 mM sodium phosphate, 20 mM sodium chloride, pH 8). EOY binding was assayed by fluorescence polarization with a synergy neo2 plate reader (*Biotek*) using a polarization filter cube (FP 485/530, *Biotek*). The binding signals were normalized to EOY4D2.1 as a positive control. The best 36 variants from libraries (200 colonies from 100 ng template library, 200 colonies from 10 ng template library) were rescreened in two 96-well plates in sextuplicates. The best variant, EOY4D2.2 2G2, was dubbed EOY4D2.2 and stemmed from the 10 ng template library. EOY4D2.2 carried only the T24M mutation, as revealed by Sanger sequencing (*Eurofins*). In cell lysate, the binding signals of EOY4D2.1 and EOY4D2.2 were 1.08-fold (p = 0.0007) and 1.14-fold (p = <0.0001) higher compared to m4D2 (n = 12, 6 and 12, respectively).

### Protein production and purification

#### Protein production

Proteins were expressed in T7 Express *E. coli* cells (*NEB*). Transformed *E. coli* colonies were grown overnight under shaking in 10 ml LB with carbenicillin (50 μg/ml). 1 ml of the overnight culture was transferred to 1L of LB, which was incubated under shaking at 37°C to an OD^600nm^ of 0.7 before induction by adding IPTG (1 mM). Proteins were expressed overnight under shaking at 18°C, after which cells were harvested by centrifugation at 4,000xg for 20 min and resuspended in assay buffer (20 mM sodium phosphate, 20 mM sodium chloride, pH 8). To lyse the cells, lysozyme and DNase I were added, and the cell suspension was frozen overnight at -20°C. Cells were thawed and further lysed with a FB-120 sonication at 4°C (1 s on, 1 s off, 3 min, 75% amplitude, 6 mm probe, *Fisherbrand*). The cell debris were removed by centrifugation at 18,000xg for 30 min.

#### Apo protein purification

Proteins were purified from the lysate by nickel affinity chromatography (5 ml HisTrap column, *GE*). After loading the lysate onto the column, the column was washed with 100 ml wash buffer (50 mM sodium phosphate, 300 mM sodium chloride, 20 mM imidazole, pH 8), and the protein was eluted with elution buffer (50 mM sodium phosphate, 300 mM sodium chloride, 250 mM imidazole, pH 8). The buffer of the eluted protein was subsequently exchanged using a G25 column (*Sigma-Aldrich*) into TEV buffer (50 mM Tris, 0.5 mM EDTA, pH 8). TEV cleavage was performed by adding TEV protease (200 μg per 1 l of culture) and TCEP (1 mM) overnight at room temperature. The cleaved designs were further purified by reverse nickel chromatography. To that end, a 5 ml HisTrap column (*GE*) was equilibrated in wash buffer (50 mM sodium phosphate, 300 mM sodium chloride, 20 mM imidazole, pH 8). Subsequently, the digestion reaction was loaded onto the column, and the flowthrough was collected. The column was washed with an additional 10 ml of wash buffer, and the flowthrough was collected again. The buffer of the combined flowthrough was exchanged into assay buffer (20 mM sodium phosphate, 20 mM sodium chloride, pH 8) using a G25 (*Sigma-Aldrich*) column. The protein samples were concentrated using centrifugal concentrators with a molecular weight cut-off of 3 kDa (*Vivaspin /Amicon*). The apo-protein was obtained after purification by size exclusion chromatography (HiLoad 16/600 Superdex 75pg, *GE*) in assay buffer (20 mM sodium phosphate, 20 mM sodium chloride, pH 8). The protein concentration was determined using the calculated extinction coefficient^39^ at 280 nm with a nanodrop. Particularly for the evolved EOY4D2.2, the apo sample was already partially loaded with cellular heme.

#### Heme-loaded protein purification

Apo protein samples were diluted to 45 ml with assay buffer (20 mM sodium phosphate, 20 mM sodium chloride, pH 8), and 1.5 equivalents of heme were added slowly to the sample (in 5 ml 20% DMSO) to load the protein with heme. The protein samples were concentrated using centrifugal concentrators with a molecular weight cut-off of 3 kDa (*Vivaspin /Amicon*). The excess heme was removed by gel filtration using size exclusion chromatography (HiLoad 16/600 Superdex 75pg, *GE*) in assay buffer (20 mM sodium phosphate, 20 mM sodium chloride, pH 8). The free heme stuck to the column during purification but could be completely removed by washing the column with 2 column volumes of water. The protein concentration was determined using the pyridine hemochrome method.^40^ To that end, the protein sample was diluted 4-10 times with 0.5 M NaOH, 10% pyridine, and absorbance spectra were measured between 500-700 nm. After reducing the sample with the addition of sodium dithionite, absorbance spectra were measured again. The heme concentration was determined from the difference in absorbance between the oxidized (540 nm) and reduced (556 nm) spectra using an extinction coefficient of 23,970 M^-1^ cm^-1^.

### Protein characterization

#### CD spectra and melt curves

Circular Dichroism (CD) spectra and melt curves were recorded using a J-1500 spectrophotometer (*JASCO*). Protein samples were prepared at a concentration of 10 μM, and the CD was baselined against assay buffer (20 mM sodium phosphate, 20 mM sodium chloride, pH 8). CD spectra were recorded between 200 nm and 250 nm at 25°C. Melt curves were recorded at 222 nm while changing the temperature between 5-95°C at a rate of 1°C per minute. Circular dichroism is reported as mean residual ellipticity (MRE) by converting the raw ellipticity θ using Eq. 1, where n is the number of peptide bonds, d is the pathlength, and c is the protein concentration.

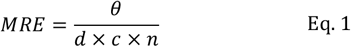

#### ITC binding assay

ITC experiments were performed with a MicroCal iTC200 (*Malvern Panalytical*). To minimize background, the ligand was dissolved in physically the same buffer batch used during the last size exclusion run during protein purification (assay buffer, 20 mM sodium phosphate, 20 mM sodium chloride, pH 8). For the EOY4D2 variants, 500 µM ligand (EOY or FLU) were titrated into 50 µM in 40 injections (200 s spacing). For the C45-based design, 1 mM EOY was titrated into 36 µM protein in 20 injections (270 s spacing).

#### Absorbance, fluorescence, and fluorescence polarization binding assay

Binding was confirmed using a variety of optical readouts using a synergy neo2 plate reader (*Biotek*). All measurements were performed in assay buffer (20 mM sodium phosphate, 20 mM sodium chloride, pH 8). For absorbance and fluorescence measurements, 1 µM EOY was titrated with varying amounts of protein. Absorbance-based binding was determined from the absorbance ratio of 538 nm to 514 nm. Fluorescence-based binding was assayed by excitation at 520 nm and emission at 540 nm. Fluorescence-polarization was assayed with a polarization filter cube (FP 485/530, *Biotek*) using 0.1 µM EOY and 1 µM enzyme.

#### CD binding assay

CD binding titrations were performed with a J-1500 spectrophotometer (*JASCO*). 800 µl enzyme (10 µM) in assay buffer (20 mM sodium phosphate, 20 mM sodium chloride, pH 8) were loaded into a 1 cm pathlength quartz cuvette. Another 500 µl enzyme (10 µM) sample was prepared containing 250 µM ligand, and the enzyme-ligand sample was added in 15 steps according to Tab. S1.

**Tab. S1:** CD titration protocol.

| Injection | V<sub>added</sub> (μl) | V<sub>total</sub> (μl) | C<sub>ligand</sub> (μM) |
| --- | --- | --- | --- |
| 1 | 0 | 0 | 0 |
| 2 | 1 | 1 | 0.3 |
| 3 | 1.5 | 2.5 | 0.8 |
| 4 | 2 | 4.5 | 1.4 |
| 5 | 3 | 7.5 | 2.3 |
| 6 | 4 | 11.5 | 3.5 |
| 7 | 6 | 17.5 | 5.4 |
| 8 | 8 | 25.5 | 7.7 |
| 9 | 10 | 35.5 | 10.6 |
| 10 | 15 | 50.5 | 14.8 |
| 11 | 20 | 70.5 | 20.2 |
| 12 | 30 | 100.5 | 27.9 |
| 13 | 40 | 140.5 | 37.3 |
| 14 | 60 | 200.5 | 50.1 |
| 15 | 80 | 280.5 | 64.9 |
| 16 | 100 | 380.5 | 80.6 |

The *K*_D_ was determined by globally fitting the CD absorbances (A) between 385 nm and 435 nm, as well as 535 nm and 560 nm, both assayed in 5 nm intervals, to Eq. 2, with the *K*_D_ shared between all wavelengths. Since a relatively high enzyme concentration [enz] = 10 µM compared to the EOY concentrations [EOY] was used, the *K*_D_ was calculated using a quadratic binding equation. Eq. 2 also contains a linear term to account for the non-specific binding (ns) as well as the fitting parameters A_max_ and c.

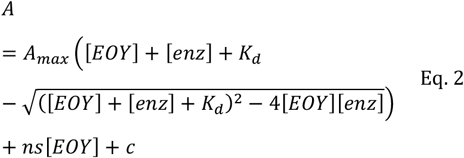

#### Mass spectrometry

The molecular weight of the protein variants was measured by electrospray ionization-mass spectrometry (ESI-MS). 10 µl of protein sample in assay buffer (20 mM sodium phosphate, 20 mM sodium chloride, pH 8) was injected onto a C4 column (*Agilent*) and eluted in a water (1% FA) / acetonitrile (1% FA) gradient (5% to 95% in 10 min, 0.25 ml/min) to remove the buffer salts. Data were acquired using a Xevo G2-XS QTof LC-MS (*Waters*) with a capillary voltage of 3.6 kV and cone voltage of 50 V in positive ion mode. The protein MS spectrum was deconvoluted using the MaxEnt-1 algorithm to obtain the actual mass of the protein. In all cases, the heme was lost during the LC/MS analysis due to the protonation of the coordinating histidines and dissociation of the bound heme under the acidic assay conditions (Tab. S2).

**Tab. S2:** Molecular weights determined by LC/MS.

| Variant | Expected mass (u) | Actual mass (u) | Difference (u) |
| --- | --- | --- | --- |
| EOY4D2 | 12236.7 | 12236.9 | 0.2 |
| EOY4D2.1 | 12735.9 | 12735.6 | -0.4 |
| EOY4D2.2 | 12765.7 | 12765.6 | -0.1 |
| EOY4D2NH | 12529.2 | 12529.5 | 0.4 |
| EOYe4D2 | 22056.3 | 22055.8 | -0.5 |
| m4D2 | 12874.6 | 12874.5 | -0.1 |
| 4D2 | 12632.0 | 12632.1 | 0.1 |
| e4D2 | 24914.0 | 24916.0 | 2.0 |

### Photo-electrochemistry

#### Redox potentiometry

Heme redox potentials were measured by optically transparent thin-layer electrochemistry (OTTLE), as described previously.^32^ In contrast to previous work, no redox mediators were added, and the equilibration time at each redox potential was thus increased to 1 h. All designed proteins were prepared at a concentration of 20 μM in assay buffer (20 mM sodium phosphate, 20 mM sodium chloride, pH 8) with 10% glycerol. If indicated, 20 μM EOY was added to the solution. UV-visible spectra were measured using a Cary 60 spectrophotometer while altering the potential using a thin platinum electrode between −225 to −525 mV vs. a silver chloride electrode, controlled by a *Biologic* SP-150 potentiostat. Potentials were adjusted vs. the Nernst hydrogen electrode (NHE) by calibration with cytochrome *c*, resulting in an average adjustment of +225 mV. Midpoint potentials (E_m_) were derived by fitting oxidation and reduction data to the Nernst equation (Eq. 3), where E is the applied potential in mV, and A_426nm_ is the absorbance of the reduced Soret peak (426 nm), and A_0_ and c are fitting parameters.

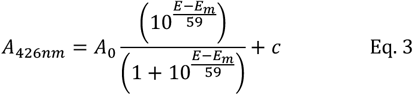

#### Photovoltaic

The photovoltaic activity of EOY, m4D2 with EOY, and EOY4D2.2 with EOY was determined using a µStat400 (*Metrohm*) and thin layer flow-cell screen-printed electrodes (TLFCL110, working electrode: carbon, auxiliary electrode: carbon, reference electrode: silver, *Metrohm*). The flow-cell electrode was filled with the respective samples and not operated in-flow. Instead, the flow cell was used to reproducibly perform the measurements with the same path length after the setup was filled with the sample. Short circuit photocurrents were measured by setting the applied potential to 0 V. Samples contained EOY (20 µM), enzyme (20 µM), and the redox mediator K_3_Fe(CN)_6_ (5 µM). Samples were excited with a green LED and mounted on a metal-core printed circuit board (M530D3, 530 nm, 480 mW, *Thorlabs*). The LED was fixed 5.8 cm above the electrode and was driven with a LEDD1B T-Cube LED Driver (*Thorlabs*) controlled by a USB-6000 controller (*Farnell*). After the potentiostat was started, the LED cycled between on (301 s) and off (301 s) with an initial delay of 601 s before switching the LED on for the first time.

Photocurrents were baselined as illustrated in Fig. S4 in the following way: First, the current at each the time the LED was switched on was determined by a linear fit of the current in the 100 s before. Next, to avoid fitting artifacts at the beginning and end of the experiment, the first and last determined currents were added five times before and after the data with a spacing of 601 s. These time points were then fitted to a polynomial, with the order of the polynomial equal to the number of points provided. The polynomial was then subtracted from the data, and the photocurrents were determined by calculating the area under the curve between each time the LED was switched on. The final photocurrent (I_final_) was determined by fitting the decaying photocurrent (I) to a single exponential function (Eq. 4). The error in the fold-change between EOY and EOY4D2.2 + EOY was determined from the standard deviation of leave-one out calculations in which one of the measured points used to calculate the fold-change was removed.

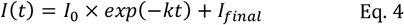

#### Potential calibration for the photovoltaic setup

The potential was calibrated against K_3_Fe(CN)_6_ using circular voltammetry against (expected potential: 436 mV, conditions: 50 mM K_3_Fe(CN)_6_ in 20 mM sodium phosphate, 20 mM sodium chloride, pH 8.0, Ebegin: 0.4V; Evtxt1: 0.4 V; Evtxt2: -0.3V; Estep: 0.002 V; scan rate: 0.01 V/s). The circular voltammetry measurement was repeated ten times, and the last three curves were used to determine the true potential.

