## Supplementary Information for "Photovoltaic enzymes by design and evolution"

### 1. Supplementary Figures

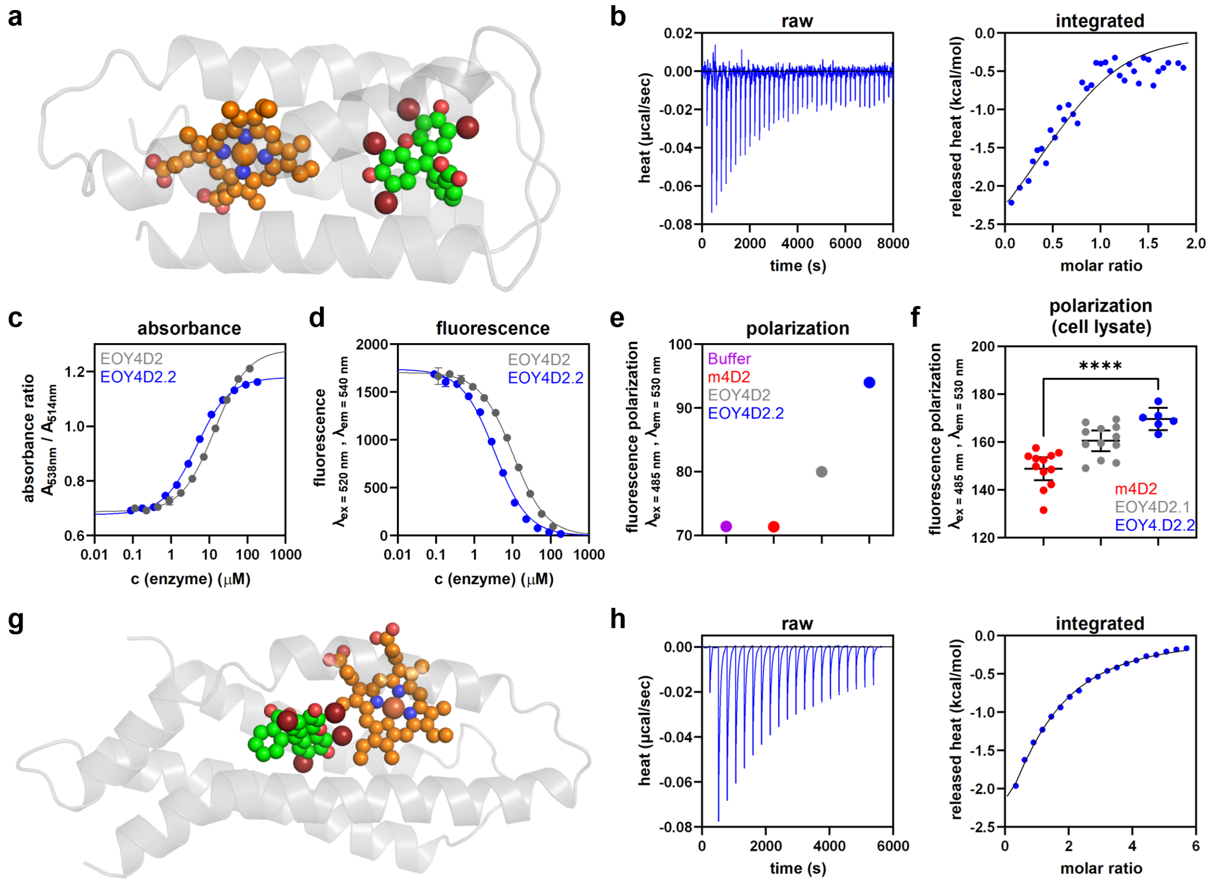


**Fig. S1 | Binding assays a,** Model of EOY4D2.2 **b,** ITC binding confirms EOY binding to EOY4D2.2. **c-e,** Absorbance, fluorescence, and fluorescence-polarization-based assays show improvements in binding during evolution (Buffer, purple; m4D2, red; EOY4D2, grey; EOY4D2.2, blue; for **c-d:** 1 µM EOY; for **e:** 0.1 µM EOY and 1 µM enzyme). **f,** Statistically significant differences in fluorescence polarization stemming from EOY binding were observed in cell lysate between EOY and EOY with either EOY4D2.1 and EOY4D2.2. **g,** Model of the C45-based design. **h,** ITC titrations confirm EOY binding to this design with a $K_d$ of 57 ± 5 µM.

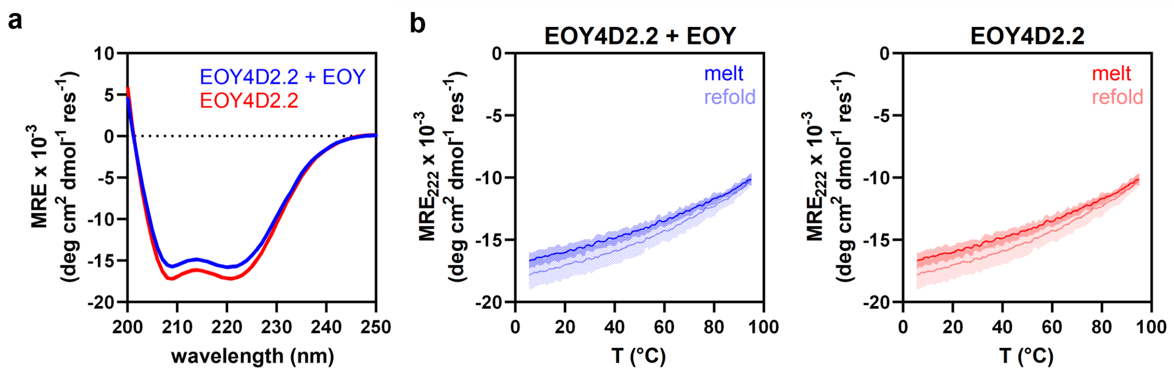


**Fig. S2 | CD spectra and melt curves of EOY4D2.2. a,** CD spectra of EOY4D2.2 with and without EOY recorded at 25°C show pronounced helicity, but are almost identical. **b,** The CD melt curves show that EOY4D2.2 is highly thermostable, as the protein is not completely unfolded at 95°C. The addition of EOY to EOY4D2.2 did not significantly affect thermal unfolding.

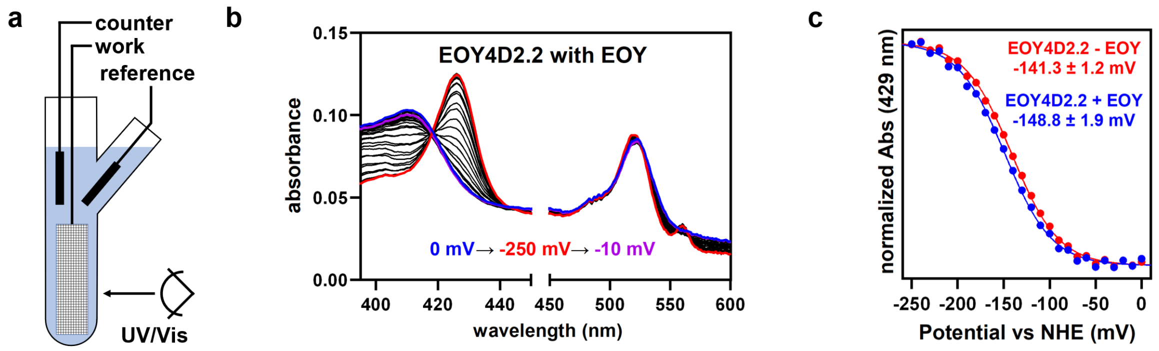


**Fig. S3 | Potentiometry. a,** Optically transparent thin-layer electrochemistry (OTTLE) setup used to assay the redox potential of EOY4D2.2 with and without EOY. **b,** Spectral changes upon cycling the redox potential from 0 mV to -250 mV to -10 mV. **c,** Fitting of the absorbance change at 429 nm reveals that the heme redox potential of EOY4D2.2 decreases from -141.3 ± 1.2 mV to -148.8 ± 1.9 mV upon EOY binding.

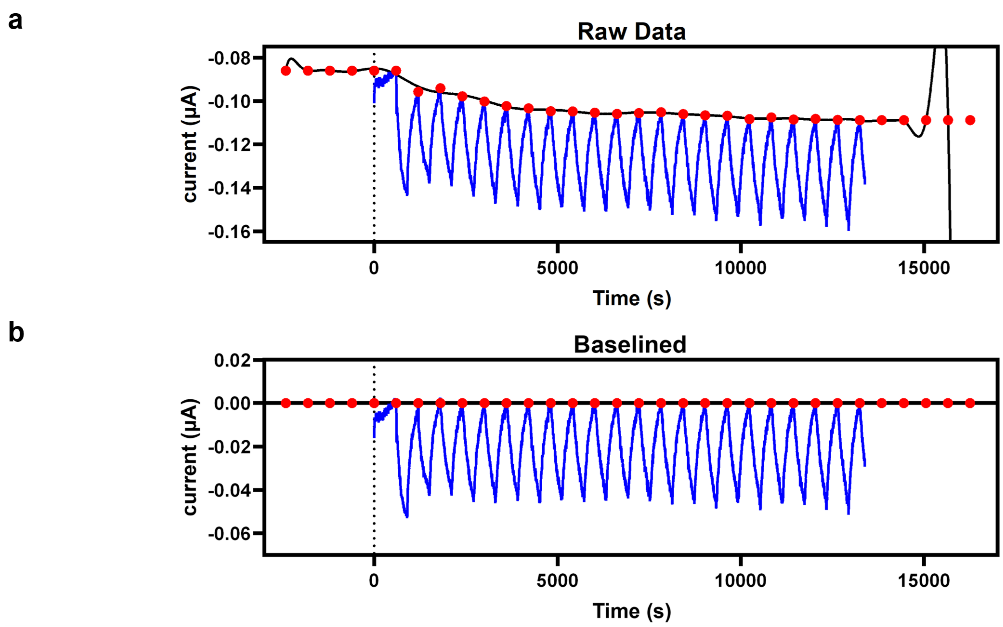


**Fig. S4 | Baselining of photovoltaic data. a,** From the raw photovoltaic data (blue), the current at each the time the LED was switched on was determined by a linear fit of the current in the 100 s before. To avoid fitting artifacts at the beginning and end of the experiment, the first and last determined currents were added five times before and after the data with a spacing of 600 s (red points). These time points were then fitted to a polynomial, with the order of the polynomial equal to the number of points provided (black line). **b,** Subtracting the baseline from (a) resulted in these baselined data.

### 2. Sequences

**DNA sequences of selected designs.** Start codons (blue), stop codons (red), $his_6$-tags (orange), and TEV cleavage sites (NLYFQ|G, purple) are highlighted.

**EOY4D2** in pET-45b(+)

ATGGCACATCATCACCATCATCATGGTAAGCCTATCCCTAACCCTCTCCTCGGTCTCGATTCTACGGAAAACCT
GTATTTTCAGGGATCGCCAGAACTGCGCGAGAAACACCGTGCGTTAGCCGAACAAGCGTACGCCGCAGCCCTAG
AAGCGATCAAGAACGAGAGCGCTTCGCCGGAACTTATCGAGAAACTTCGTGCTGCGACAGAACAGGTGTATGCG
ACTGGCCAGGAAATGCTGAAAAACGGGTCTGTAAGTCCGTCACCTGAACTGCGGGAGAAACACCGCGCTTTGGC
CGAACAGGCTTACGCAGCCGCTGCGGAGCAGCGCCAGACCAAGTCCACTAGCCCGGAAAAGATTGAGAAAGCTG
CCGCATTACAGGAACAAGTCTATGCGACCGGTCAGGAAATGTTGAAAAATTGA

**EOY4D2.1** in pET-21(+)

ATGCATCATCATCACCACCACGGCAAACCTATTCCTAACCCGCTGCTGGGCCTGGATAGCACGGAAAACTTATA
TTTTCAGGGTAGCCCCGAACTCCGTGAGAAACACCGCGCTCTGGCGGAACAGGCGTATGCGATTGCACTGGAGA
TGACGAAAAATGAATCATGGTCACCTGAATTGGTGGAAAAGCTCCGTGCCATCTTGGAACAGGTCTATGCGACT
GGGCAAGAGATGCTGAAAAATGGCTCCGTCTCCCCTAGCCCTGAATTGCGCGAAAAACACCGTGCACTGGCTGA
GCAGCTGTACGCCTATGCTGCTATTCAAAAACAAACTGTTTTTACCAGCCCAGAAAAAAAAGAGAAAATGAAGG
CGCTGCAAGAACAGGTTTATGCCACCGGGCAAGAAATGCTGAAAAACTAA

**EOY4D2.2** in pET-21(+)

ATGCATCATCATCACCACCACGGCAAACCTATTCCTAACCCGCTGCTGGGCCTGGATAGCACGGAAAACTTATA
TTTTCAGGGTAGCCCCGAACTCCGTGAGAAACACCGCGCTCTGGCGGAACAGGCGTATGCGATTGCACTGGAGA
TGATGAAAAATGAATCATGGTCACCTGAATTGGTGGAAAAGCTCCGTGCCATCTTGGAACAGGTCTATGCGACT
GGGCAAGAGATGCTGAAAAATGGCTCCGTCTCCCCTAGCCCTGAATTGCGCGAAAAACACCGTGCCCTGGCTGA
GCAGCTGTACGCCTATGCTGCTATTCAAAAACAAACTGTTTTTACCAGCCCAGAAAAAAAAGAGAAAATGAAGG
CGCTGCAAGAACAGGTTTATGCCACCGGTCAAGAAATGCTGAAAAACTAA

**EOY4D2NH** in pET-45b(+)

ATGGCACATCATCACCATCATCATGGTAAGCCTATCCCTAACCCTCTCCTCGGTCTCGATTCTACGGAAAACCT
GTATTTTCAGGGATCGCCAGAACTGCGCGAGAAACTCCGTGCGTTAATCGAACAAGCGTACGCCGCAGCCCTAG
AAGCGATCAAGAACGAGAGCGCTTCGCCGGAACTTATCGAGAAACTTCGTGCTGCGACAGAACAGGTGTATGCG
ACTTGGCAGGAATTACTGAAAAACGGGTCTGTAAGTCCGTCACCTGAACTGCGGGAGAAATTCCGCGCTTTGCT
CGAACAGGCTTACGCAGCCGCTGCGGAGCAGCGCCAGACCAAGTCCACTAGCCCGGAAAAGATTGAGAAAGCTG
CCGCATTACAGGAACAAGTCTATGCGACCTGGCAGGAACTGTTGAAAAATTGA

**EOYe4D2** in pET-21(+)

ATGCACCATCACCACCATCACGGCAAGCCAATTCCGAACCCTCTGCTTGGCCTTGATTCCACCGAAAATTTGTA
CTTCCAGGGTTCGCCAGAACTTCGCGAGAAGCACCGCGCCCTCGCCGAGCAAGTGTATGCGACAGGGCAAGAGA
TGCTCGAGCTGCGTGAAAAGCATCGCGCCTTGGCGGAGCAGGCGTATGCGGCAGCACTTGAAGCGATTAAGAAC
GAGTCGGCGTCCCCAGAGTTAATTGAGAAGCTCCGCGCGGCTACTGAGCAGGTTTACGCTACAGGTCAGGAGAT
GTTGGAGCTTCGTGAAAAGCATCGCGCTTTGGCGGAACAGGTTTATGCCACGGGGCAAGAGATGCTTAAGAATG
GCAGTGTGTCACCGAGTCCAGAACTGCGTGAGAAACATCGTGCTCTTGCGGAGCAGGTTTATGCCACCGGCCAG
GAAATGTTAGAGTTGCGCGAGAAGCATCGTGCACTGGCCGAGCAGGCGTATGCAGCCGCCGCCGAACAGCGTCA
AACAAAATCCACCAGTCCGGAGAAAATTGAAAAGGCGGCTGCACTGCAGGAACAAGTATACGCCACTGGCCAAG
AAATGCTGGAGCTGCGTGAAAAACACCGCGCCCTGGCAGAACAGGTGTACGCTACCGGACAAGAAATGTTGAAA
AATTAA

**EOY4D2.2:** Full sequence of this variant in pET-21(+)

```
TTAACTTTAAGAAGGAGATATACATATGCATCATCATCACCACCACGGCAAACCTATTCCTAACCCGCTGCTGGGCCTGGATAGCACGGAAAACTTATATTTTCAGGGTAGCCCCGAACTCCGTGAGAAACACCGCGCTCTGGCGGAACAGGCGTATGCGATTGCACTGGAGATGATGAAAAATGAA
TCATGGTCACCTGAATTGGTGGAAAAGCTCCGTGCCATCTTGGAACAGGTCTATGCGACTGGGCAAGAGATGCTGAAAAATGGCTCCGTCTCCCCTAGCCCTGAATTGCGCGAAAAACACCGTGCCCTGGCTGAGCAGCTGTACGCCTATGCTGCTATTCAAAAACAAACTGTTTTTACCAGCCCAG
AAAAAAAAGAGAAAATGAAGGCGCTGCAAGAACAGGTTTATGCCACCGGTCAAGAAATGCTGAAAAACTAACTCGAGCACCACCACCACCACCACTGAGATCCGGCTGCTAACAAAGCCCGAAAGGAAGCTGAGTTGGCTGCTGCCACCGCTGAGCAATAACTAGCATAACCCCTTGGGGCCTCTAA
ACGGGTCTTGAGGGGTTTTTTGCTGAAAGGAGGAACTATATCCGGATTGGCGAATGGGACGCGCCCTGTAGCGGCGCATTAAGCGCGGCGGGTGTGGTGGTTACGCGCAGCGTGACCGCTACACTTGCCAGCGCCCTAGCGCCCGCTCCTTTCGCTTTCTTCCCTTCCTTTCTCGCCACGTTCGCCG
GCTTTCCCCGTCAAGCTCTAAATCGGGGGCTCCCTTTAGGGTTCCGATTTAGTGCTTTACGGCACCTCGACCCCAAAAAACTTGATTAGGGTGATGGTTCACGTAGTGGGCCATCGCCCTGATAGACGGTTTTTCGCCCTTTGACGTTGGAGTCCACGTTCTTTAATAGTGGACTCTTGTTCCAAAC
TGGAACAACACTCAACCCTATCTCGGTCTATTCTTTTGATTTATAAGGGATTTTGCCGATTTCGGCCTATTGGTTAAAAAATGAGCTGATTTAACAAAAATTTAACGCGAATTTTAACAAAATATTAACGCTTACAATTTAGGTGGCACTTTTCGGGGAAATGTGCGCGGAACCCCTATTTGTTTAT
TTTTCTAAATACATTCAAATATGTATCCGCTCATGAGACAATAACCCTGATAAATGCTTCAATAATATTGAAAAAGGAAGAGTATGAGTATTCAACATTTCCGTGTCGCCCTTATTCCCTTTTTTGCGGCATTTTGCCTTCCTGTTTTTGCTCACCCAGAAACGCTGGTGAAAGTAAAAGATGCTGA
AGATCAGTTGGGTGCACGAGTGGGTTACATCGAACTGGATCTCAACAGCGGTAAGATCCTTGAGAGTTTTCGCCCCGAAGAACGTTTTCCAATGATGAGCACTTTTAAAGTTCTGCTATGTGGCGCGGTATTATCCCGTATTGACGCCGGGCAAGAGCAACTCGGTCGCCGCATACACTATTCTCAG
AATGACTTGGTTGAGTACTCACCAGTCACAGAAAAGCATCTTACGGATGGCATGACAGTAAGAGAATTATGCAGTGCTGCCATAACCATGAGTGATAACACTGCGGCCAACTTACTTCTGACAACGATCGGAGGACCGAAGGAGCTAACCGCTTTTTTGCACAACATGGGGGATCATGTAACTCGCC
TTGATCGTTGGGAACCGGAGCTGAATGAAGCCATACCAAACGACGAGCGTGACACCACGATGCCTGCAGCAATGGCAACAACGTTGCGCAAACTATTAACTGGCGAACTACTTACTCTAGCTTCCCGGCAACAATTAATAGACTGGATGGAGGCGGATAAAGTTGCAGGACCACTTCTGCGCTCGGC
CCTTCCGGCTGGCTGGTTTATTGCTGATAAATCTGGAGCCGGTGAGCGTGGGTCTCGCGGTATCATTGCAGCACTGGGGCCAGATGGTAAGCCCTCCCGTATCGTAGTTATCTACACGACGGGGAGTCAGGCAACTATGGATGAACGAAATAGACAGATCGCTGAGATAGGTGCCTCACTGATTAAG
CATTGGTAACTGTCAGACCAAGTTTACTCATATATACTTTAGATTGATTTAAAACTTCATTTTTAATTTAAAAGGATCTAGGTGAAGATCCTTTTTGATAATCTCATGACCAAAATCCCTTAACGTGAGTTTTCGTTCCACTGAGCGTCAGACCCCGTAGAAAAGATCAAAGGATCTTCTTGAGATC
CTTTTTTTCTGCGCGTAATCTGCTGCTTGCAAACAAAAAAACCACCGCTACCAGCGGTGGTTTGTTTGCCGGATCAAGAGCTACCAACTCTTTTTCCGAAGGTAACTGGCTTCAGCAGAGCGCAGATACCAAATACTGTCCTTCTAGTGTAGCCGTAGTTAGGCCACCACTTCAAGAACTCTGTAGC
ACCGCCTACATACCTCGCTCTGCTAATCCTGTTACCAGTGGCTGCTGCCAGTGGCGATAAGTCGTGTCTTACCGGGTTGGACTCAAGACGATAGTTACCGGATAAGGCGCAGCGGTCGGGCTGAACGGGGGGTTCGTGCACACAGCCCAGCTTGGAGCGAACGACCTACACCGAACTGAGATACCTA
CAGCGTGAGCTATGAGAAAGCGCCACGCTTCCCGAAGGGAGAAAGGCGGACAGGTATCCGGTAAGCGGCAGGGTCGGAACAGGAGAGCGCACGAGGGAGCTTCCAGGGGGAAACGCCTGGTATCTTTATAGTCCTGTCGGGTTTCGCCACCTCTGACTTGAGCGTCGATTTTTGTGATGCTCGTCAG
GGGGGCGGAGCCTATGGAAAAACGCCAGCAACGCGGCCTTTTTACGGTTCCTGGCCTTTTGCTGGCCTTTTGCTCACATGTTCTTTCCTGCGTTATCCCCTGATTCTGTGGATAACCGTATTACCGCCTTTGAGTGAGCTGATACCGCTCGCCGCAGCCGAACGACCGAGCGCAGCGAGTCAGTGAG
CGAGGAAGCGGAAGAGCGCCTGATGCGGTATTTTCTCCTTACGCATCTGTGCGGTATTTCACACCGCAATGGTGCACTCTCAGTACAATCTGCTCTGATGCCGCATAGTTAAGCCAGTATACACTCCGCTATCGCTACGTGACTGGGTCATGGCTGCGCCCCGACACCCGCCAACACCCGCTGACGC
GCCCTGACGGGCTTGTCTGCTCCCGGCATCCGCTTACAGACAAGCTGTGACCGTCTCCGGGAGCTGCATGTGTCAGAGGTTTTCACCGTCATCACCGAAACGCGCGAGGCAGCTGCGGTAAAGCTCATCAGCGTGGTCGTGAAGCGATTCACAGATGTCTGCCTGTTCATCCGCGTCCAGCTCGTTG
AGTTTCTCCAGAAGCGTTAATGTCTGGCTTCTGATAAAGCGGGCCATGTTAAGGGCGGTTTTTTCCTGTTTGGTCACTGATGCCTCCGTGTAAGGGGGATTTCTGTTCATGGGGGTAATGATACCGATGAAACGAGAGAGGATGCTCACGATACGGGTTACTGATGATGAACATGCCCGGTTACTGG
AACGTTGTGAGGGTAAACAACTGGCGGTATGGATGCGGCGGGACCAGAGAAAAATCACTCAGGGTCAATGCCAGCGCTTCGTTAATACAGATGTAGGTGTTCCACAGGGTAGCCAGCAGCATCCTGCGATGCAGATCCGGAACATAATGGTGCAGGGCGCTGACTTCCGCGTTTCCAGACTTTACGA
AACACGGAAACCGAAGACCATTCATGTTGTTGCTCAGGTCGCAGACGTTTTGCAGCAGCAGTCGCTTCACGTTCGCTCGCGTATCGGTGATTCATTCTGCTAACCAGTAAGGCAACCCCGCCAGCCTAGCCGGGTCCTCAACGACAGGAGCACGATCATGCGCACCCGTGGGGCCGCCATGCCGGCG
ATAATGGCCTGCTTCTCGCCGAAACGTTTGGTGGCGGGACCAGTGACGAAGGCTTGAGCGAGGGCGTGCAAGATTCCGAATACCGCAAGCGACAGGCCGATCATCGTCGCGCTCCAGCGAAAGCGGTCCTCGCCGAAAATGACCCAGAGCGCTGCCGGCACCTGTCCTACGAGTTGCATGATAAAGA
AGACAGTCATAAGTGCGGCGACGATAGTCATGCCCCGCGCCCACCGGAAGGAGCTGACTGGGTTGAAGGCTCTCAAGGGCATCGGTCGAGATCCCGGTGCCTAATGAGTGAGCTAACTTACATTAATTGCGTTGCGCTCACTGCCCGCTTTCCAGTCGGGAAACCTGTCGTGCCAGCTGCATTAATG
AATCGGCCAACGCGCGGGGAGAGGCGGTTTGCGTATTGGGCGCCAGGGTGGTTTTTCTTTTCACCAGTGAGACGGGCAACAGCTGATTGCCCTTCACCGCCTGGCCCTGAGAGAGTTGCAGCAAGCGGTCCACGCTGGTTTGCCCCAGCAGGCGAAAATCCTGTTTGATGGTGGTTAACGGCGGGAT
ATAACATGAGCTGTCTTCGGTATCGTCGTATCCCACTACCGAGATATCCGCACCAACGCGCAGCCCGGACTCGGTAATGGCGCGCATTGCGCCCAGCGCCATCTGATCGTTGGCAACCAGCATCGCAGTGGGAACGATGCCCTCATTCAGCATTTGCATGGTTTGTTGAAAACCGGACATGGCACTC
CAGTCGCCTTCCCGTTCCGCTATCGGCTGAATTTGATTGCGAGTGAGATATTTATGCCAGCCAGCCAGACGCAGACGCGCCGAGACAGAACTTAATGGGCCCGCTAACAGCGCGATTTGCTGGTGACCCAATGCGACCAGATGCTCCACGCCCAGTCGCGTACCGTCTTCATGGGAGAAAATAATAC
TGTTGATGGGTGTCTGGTCAGAGACATCAAGAAATAACGCCGGAACATTAGTGCAGGCAGCTTCCACAGCAATGGCATCCTGGTCATCCAGCGGATAGTTAATGATCAGCCCACTGACGCGTTGCGCGAGAAGATTGTGCACCGCCGCTTTACAGGCTTCGACGCCGCTTCGTTCTACCATCGACAC
CACCACGCTGGCACCCAGTTGATCGGCGCGAGATTTAATCGCCGCGACAATTTGCGACGGCGCGTGCAGGGCCAGACTGGAGGTGGCAACGCCAATCAGCAACGACTGTTTGCCCGCCAGTTGTTGTGCCACGCGGTTGGGAATGTAATTCAGCTCCGCCATCGCCGCTTCCACTTTTTCCCGCGTT
TTCGCAGAAACGTGGCTGGCCTGGTTCACCACGCGGGAAACGGTCTGATAAGAGACACCGGCATACTCTGCGACATCGTATAACGTTACTGGTTTCACATTCACCACCCTGAATTGACTCTCTTCCGGGCGCTATCATGCCATACCGCGAAAGGTTTTGCGCCATTCGATGGTGTCCGGGATCTCGA
CGCTCTCCCTTATGCGACTCCTGCATTAGGAAGCAGCCCAGTAGTAGGTTGAGGCCGTTGAGCACCGCCGCCGCAAGGAATGGTGCATGCAAGGAGATGGCGCCCAACAGTCCCCCGGCCACGGGGCCTGCCACCATACCCACGCCGAAACAAGCGCTCATGAGCCCGAAGTGGCGAGCCCGATCTT
CCCCATCGGTGATGTCGGCGATATAGGCGCCAGCAACCGCACCTGTGGCGCCGGTGATGCCGGCCACGATGCGTCCGGCGTAGAGGATCGAGATCTCGATCCCGCGAAATTAATACGACTCACTATAGGGGAATTGTGAGCGGATAACAATTCCCCTCTAGGATCC
```

**Protein sequences of selected designs.** Compared to m4D2, 27 mutations were introduced in EOY4D2 (red). The no-heme variant of EOY4D2 (EOY4D2_NH, orange) carries eight additional mutations to knock out heme binding. Sixteen mutations were added in EOY4D2.1 (purple), and the T24M mutation was discovered during the evolution of EOY4D2.2 (blue).

```
            1          11         21         31         41
4D2         GSPELREKHR ALAEQVYATG QEMLKNTSNS PELREKHRAL AEQVYATGQE
EOY4D2      GSPELREKHR ALAEQAYAAA LEAIKNESAS PELIEKLRAA TEQVYATGQE
EOY4D2_NH   GSPELREKLR ALIEQAYAAA LEAIKNESAS PELIEKLRAA TEQVYATWQE
EOY4D2.1    GSPELREKHR ALAEQAYAIA LEMTKNESWS PELVEKLRAI LEQVYATGQE
EOY4D2.2    GSPELREKHR ALAEQAYAIA LEMMKNESWS PELVEKLRAI LEQVYATGQE

            51         61         71         81         91
4D2         MLKNGSVSPS PELREKHRAL AEQVYATGQE MLKNTSNSPE LREKHRALAE
EOY4D2      MLKNGSVSPS PELREKHRAL AEQAYAAAAE QRQTKSTSPE KIEKAAALQE
EOY4D2_NH   LLKNGSVSPS PELREKFRAL LEQAYAAAAE QRQTKSTSPE KIEKAAALQE
EOY4D2.1    MLKNGSVSPS PELREKHRAL AEQLYAYAAI QKQTVFTSPE KKEKMKALQE
EOY4D2.2    MLKNGSVSPS PELREKHRAL AEQLYAYAAI QKQTVWTSPE KKEKMKALQE

            101        111
4D2         QVYATGQEML KN*
EOY4D2      QVYATGQEML KN*
EOY4D2_NH   QVYATWQELL KN*
EOY4D2.1    QVYATGQEML KN*
EOY4D2.2    QVYATGQEML KN*
```

Compared to C45, 12 mutations were introduced during computational design resulting in the C45-based design EOY342 (blue) binding EOY with a $K_d$ of 57 ± 5 µM.

```
            1          11         21         31         41
C45         GMTPEQIWKQ FEDALQKFEE ALNQFEDLKQ LGGSGSGSGG EIWKQFEDAL
EOY342      GMTPEQIWKQ FEDAAQKAEE ALNQFEDLKQ LGGSGSGSGG EIWKQFEDAL

            51         61         71         81         91
C45         QKFEEALNQF EDLKQLGGSG GSGGSGGEIW KQFEDALQKF EEALNQFEDL
EOY342      QKKAEKAKQF EDLKQLGGSG GSGGSGGEIW KQFEDALQKA LEALNQFEDL

            101        111        121        131
C45         KQLGGSGSGS GGECIACHED ALQKFEEALN QFEDLKQL*
EOY342      KQLGGSGSGS GGECIACHED ALQKAEEATN DFEDLKQL*
```
